# Bacteria defend against phages through type IV pilus glycosylation

**DOI:** 10.1101/150227

**Authors:** Hanjeong Harvey, Joseph Bondy-Denomy, Hélène Marquis, Kristina M. Sztanko, Alan R. Davidson, Lori L. Burrows

**Author notes:** For correspondence: Lori L. Burrows:, Alan R. Davidson.

## Abstract

Bacterial surface structures such as type IV pili are common receptors for phage. Strains of the opportunistic pathogen *Pseudomonas aeruginosa* express one of five different major type IV pilin alleles, two of which are glycosylated with either lipopolysaccharide O-antigen units or polymers of D-arabinofuranose. Here we show that both these post-translational modifications protect *P. aeruginosa* from a variety of pilus-specific phages. We identified a phage capable of infecting strains expressing both non-glycosylated and glycosylated pilins, and through construction of a chimeric phage, traced this ability to its unique tail proteins. Alteration of pilin sequence, or masking of binding sites by glycosylation, both block phage infection. The energy invested by prokaryotes in glycosylating thousands of pilin subunits is thus explained by the protection against phage predation provided by these common decorations.

**SIGNIFICANCE:** Post-translational modification of bacterial and archaeal surface structures such as pili and flagella is widespread, but the function of these decorations is not clear. We propose that predation by bacteriophages that use these structures as receptors selects for strains that mask potential phage binding sites using glycosylation. Phages are of significant interest as alternative treatments for antibiotic-resistant pathogens, but the ways in which phage interact with host receptors are not well understood. We show that specific phage tail proteins allow for infection of strains with glycosylated pili, providing a foundation for the creation of designer phages that can circumvent first-line bacterial defenses.

## INTRODUCTION

Phages present a major challenge to bacterial survival in most environments^1^. For this reason, bacteria have evolved a battery of anti-phage defenses including CRISPR-Cas, restriction-modification, and abortive infection systems ^2, 3, 4^. Such strategies are effective at inhibiting diverse types of phage, but the phage genome – which encodes many potentially dangerous gene products – is still allowed to enter the cell. The safest and most effective way to inhibit phage infection is to block it at its initial step, phage particle adsorption to the cell surface. Here we describe a cell-surface modification of *Pseudomonas aeruginosa* that can block infection by many different phages.

Although a large number of diverse phages infecting *P. aeruginosa* have been described, they use predominantly two cell surface receptors, lipopolysaccharides (LPS) and type IV pili (T4P)^5, 6^. It is likely that the evolution of different LPS O-antigen serotypes in *P. aeruginosa* strains has been shaped by the battle with phages, since individual phage types generally infect only limited subsets of these serotypes. However, it is unclear how targeting of T4P by phages has affected the evolution of their components.

T4P are used by a wide variety of bacterial and archaeal species for adherence, biofilm formation, DNA uptake, and a form of surface-associated motility called ‘twitching’ ^7, 8, 9^. T4P are also virulence factors for many bacterial pathogens, including *P. aeruginosa* ^10, 11, 12, 13, 14^. Although many *P. aeruginosa* phages use T4P as receptors ^6, 15, 16, 17^, the exact manner in which they interact with pili remains unknown. T4P are composed of thousands of copies of the major pilin, PilA, but also contain small amounts of minor (low-abundance) pilins that are thought to prime assembly by forming an initiation complex to which major pilins are subsequently added, extending the pilus from its base in the inner membrane^18^. Thus, minor pilins are most likely positioned at the tip of assembled pili^19^, though rare incorporation along the filament may also occur ^20^. Detailed electron microscopy studies of *P. aeruginosa* pilus-specific phages suggested that they bind along the length of the pilus, and are pulled closer to the cell surface when the pilus retracts ^21^. These observations imply that the phages bind to the major pilin directly, though this assumption has not been formally tested.

Each *P. aeruginosa* strain encodes one of 5 different major pilin alleles at a conserved locus located between the *pilB* gene, encoding the pilus extension ATPase, and a tRNA^Thr^ gene^22^. The pilin variants differ with respect to their length (150 to 173 residues), the number of amino acids (12 to 29 residues) between the two C-terminal Cys residues that form a structurally and functionally critical disulfide-bonded loop or ‘D-region’, and the presence of specific pilin accessory genes immediately downstream of the *pilA* gene. A census of nearly 300 strains^22^ revealed that the most common allele is group I, a pilin that is glycosylated at the C-terminal Ser by its associated glycosyltransferase, TfpO ^23^. The pilin glycan is an O-antigen unit, sourced from the O-specific antigen (OSA, formerly B-band) lipopolysaccharide (LPS) biosynthetic pathway ^24^. The group I allele was over-represented (close to 70%) in strains isolated from cystic fibrosis patients, suggesting that it provides a significant advantage in certain environments^22^. We subsequently described a second, distinct glycosylation system in group IV strains, such as PA7 ^25, 26, 27^. The pilin subunits in this group are glycosylated by their cognate glycosyltransferase TfpW on multiple Ser and Thr residues with homopolymers of α1,5-linked D-arabinofuranose (D-Ara*f*). The genes encoding the biosynthesis of D-Araf polymers are unlinked to the pilin locus, but glycosylation is important for stable pilin expression ^26^.

A convincing reason for the prevalence of pilin glycosylation in nature has not yet been put forward, though this modification has a modest effect on pathogenicity in mice ^28^ and decreases twitching motility on plastic ^29, 30^. In *Neisseria meningitidis*, an obligate human commensal, and in the multidrug resistant genus *Acinetobacter*, pilin glycosylation was proposed to block binding of pilin–specific antibodies ^31, 32^. Here we show that modification of *P. aeruginosa* pilins by either of its glycosylation systems blocks infection by many pilus-specific phages. We further show that phages producing unique tail proteins can partially breach this defense. The ability of pilin glycosylation to protect against phage predation provides a compelling rationale for the widespread occurrence of this prokaryotic post-translational modification.

## RESULTS

### Pilin glycosylation blocks phage infection

A previous study of *P. aeruginosa* strain 1244, which expresses glycosylated group I pilins, concluded that this post-translational modification had no effect on phage susceptibility ^28^. However, that study used a phage that was capable of infecting the wild type strain, which expresses glycosylated pilins. While investigating the host range of a large number of pilus-specific phages that we isolated previously ^17^, we noted that some were capable of infecting a 1244 *tfpO* mutant, which produces unmodified pilins, but not its wild type parent. These results hinted that pilin glycosylation might represent a mechanism of resistance to phages that use the pilus as a receptor.

Since most of the phages in our collection grow poorly on the 1244 wild type or *tfpO* mutant strains, we performed further studies using the common laboratory strain PAO1, which is sensitive to many more phages. To compare the ability of different *pilA* alleles in a common genetic background to allow phage infection, we used a set of recombinant strains expressing the *pilA* genes of interest from an arabinose-inducible plasmid in a PAO1 *pilA* null mutant ^29^. We then used standard phage spotting assays ^17^ to assess the ability of 19 different T4P-dependent non-contractile tailed phages to replicate on strains expressing the native PAO1 PilA_II_ (group II), or the strain 1244 PilA_I_ (group I) in the presence or absence of TfpO, which mediates pilin glycosylation. We found that while all these phages plated robustly on the strains expressing PilA_II_ or unglycosylated PilA_I_, TfpO-mediated glycosylation of PilA_I_ with O-antigen units completely inhibited the growth of all except phage DMS3, which was partially inhibited (**Table 1**).

**Table 1.** Titer of phages on recombinant PAO1 strains

| Phages | PAO1<sup>a</sup> | PAO1 *pilA* |  |  |
| --- | --- | --- | --- | --- |
|  |  | + *pilA*<sub>I</sub><sup>b</sup> | + *pilA*<sub>I</sub>-*tfpO*<sup>c</sup> | + *pilA*<sub>II</sub><sup>d</sup> |
| DMS3 | -5 | -4 | -3 | -4 |
| JBD26 | -6 | -5 | 0 | -5 |
| JBD68 | -6 | -5 | 0 | -6 |
| JBD8 | -6 | -6 | 0 | -6 |
| JBD23 | -6 | -4 | 0 | -4 |
| JBD24 | -6 | -6 | 0 | -6 |
| JBD35c | -6 | -5 | 0 | -5 |
| JBD69a | -6 | -6 | 0 | -6 |
| MP22 | -5 | -4 | 0 | -4 |
| MP29 | -5 | -4 | 0 | -4 |
| JBD88a | -6 | -6 | 0 | -6 |
| JBD16c | -6 | -4 | 0 | -4 |
| JBD18 | -6 | -6 | 0 | -6 |
| JBD25 | -6 | -6 | 0 | -6 |
| JBD63 | -6 | -5 | 0 | -5 |
| JBD67b | -6 | -6 | 0 | -6 |
| D3112 | -6 | -5 | 0 | -6 |
| JBD5 | -6 | -5 | 0 | -5 |
| JBD10 | -6 | -4 | 0 | -4 |
| JBD30 | -6 | -6 | 0 | -6 |
| JBD93a | -5 | -4 | 0 | -5 |
| M6 | -6 | 0 | 0 | -6 |

<sup>a</sup>PAO1 wild type containing the vector pBADGr
<sup>b</sup>PAO1 *pilA* mutant expressing *pilA*<sub>I</sub> from *P. aeruginosa* 1244
<sup>c</sup>PAO1 *pilA* mutant expressing *pilA*<sub>I</sub> and *tfpO* from *P. aeruginosa*
<sup>d</sup>PAO1 *pilA* mutant expressing *pilA*<sub>II</sub> from *P. aeruginosa* PAO1

Most of the phages tested were similar in sequence and genome structure ^17^, particularly within the phage tail operon expected to be involved in host receptor binding. For this reason, we focused further studies on three phages, JBD26, DMS3, and JBD68. JBD26 and DMS3 are very similar in sequence and represent the dominant group in this collection (*P. aeruginosa* phage MP22-like) while JBD68 resembles *P. aeruginosa* phage F10 (**Supplementary Table S1**) and its putative tail proteins display little similarity to those of the other phages ^17^. All three of these phages plated robustly on a PAO1 *pilA* mutant expressing PilAI or PilAII (**Fig. 1A**). However, co-expression of PilAI and TfpO caused a greater than 10^5^-fold reduction of JBD26 and JBD68 infectivity, while infectivity of DMS3 was reduced by ~10^3^-fold (**Fig. 1A**). To further investigate this phenomenon, we mutated the C-terminal Ser of PilAI to Ala in the PilAI TfpO expressing strain, precluding attachment of the glycan to the pilin ^23^. The resulting strain produced non-glycosylated pilins and was susceptible to all three phages (**Fig. 1A**), confirming that pilin modification was responsible for resistance.

**Figure 1.**
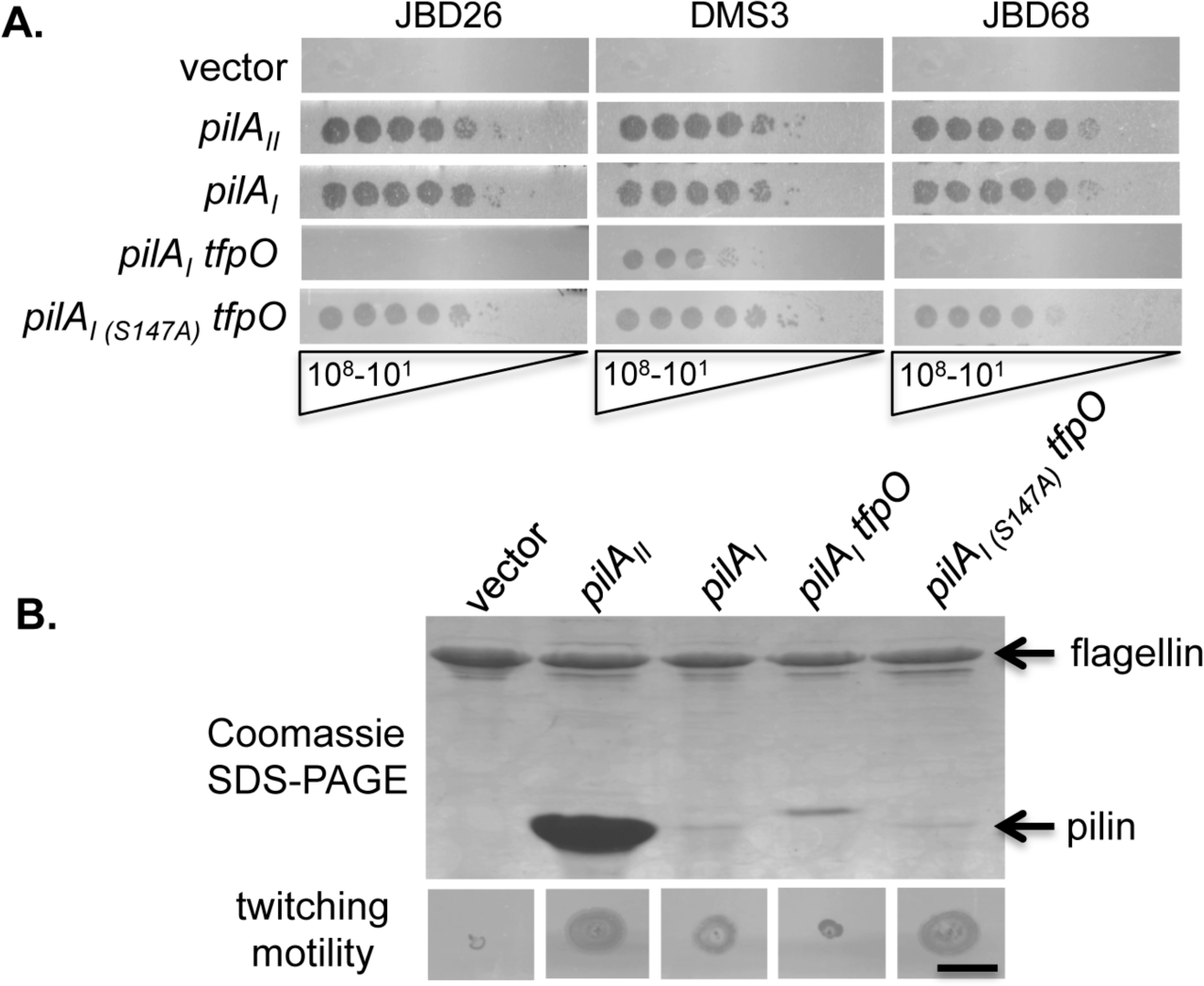
Infection of a *P. aeruginosa* PAO1 *pilA* mutant expressing non-glycosylated versus glycosylated pilins. **A.** Five μl each of serial ten-fold dilutions (10e8 to 10e1) of phages JBD26, DMS3, and JBD68 were spotted onto a PAO1 *pilA* mutant expressing the genes indicated on the left, expressed from the pBADGr vector. The top row is the empty vector control. Loss of pilin glycosylation in the S147A mutant restores phage susceptibility. **B.** Surface piliation of the PAO1 *pilA* mutant complemented with its cognate pilin (*pilA_II_*), or that of group I strain 1244 (*pilA_I_*). Differences in the levels of recoverable surface pili do not correlate with phage susceptibility or twitching motility on polystyrene. Scale bar = 1 cm.

To ensure that changes to phage susceptibility were due to changes in pilin glycosylation status, and not to differences in levels of pili on the cell surface, we examined the amount of surface pili sheared from the recombinant strains. As we showed previously ^29^, a PAO1 *pilA* mutant expressing group I pilins, regardless of their glycosylation status, assembled fewer surface pili compared to the strain expressing the native PilA_II_ and had reduced twitching motility (**Fig. 1B**). This difference in surface piliation did not affect phage susceptibility (compare *pilA_II_* versus *pilA_I_* in **Fig 1A**). Since the PilA_I_ TfpO strain expressed more surface pili than the PilAI strain expressing unmodified pilins, the reduction in plaquing efficiency on the former was not due to inadequate piliation (**Fig. 1B**).

### A putative DMS3 tail protein confers the ability to utilize glycosylated pili

Phage DMS3 was distinct from the other phages tested, in that it could replicate to some extent on cells expressing glycosylated pili. To understand what features of DMS3 might explain this ability, we compared its genome to that of JBD26, a closely related phage. Both are temperate double-stranded DNA phages with non-contractile tails. The two phages have similar genome sizes (~37 kb) and share 81% identity at the nucleotide level (Genbank accession JN811560 for JBD26, and NC_008717 for DMS3; ^33^). However, one striking difference between these phages occurs directly downstream of genes encoding their tail components. In this region, the genome of JBD26 contains two open reading frames (ORFs) – *59* and *60* – positioned between conserved ORFs *58* and *61*, while DMS3 has 3 ORFs – *49-51* – at the same position (**Fig. 2A**). Comparison of the amino acid sequences of the predicted gene products (gp) revealed that JBD26 gp59 and DMS3 gp49 are 86% identical over the first 153 residues (shown in white in **Fig 2A**), but then their sequences diverge completely (**Supplementary Fig. S1A**). Similarly, the last 47 residues of JBD26 gp60 and DMS3 gp51 are 96% identical, while the N-termini of those proteins differ. DMS3 gp50 has no counterpart in JBD26. Given the positioning of the genes encoding these proteins at the 3’-end of the tail encoding regions within their respective genomes, we hypothesized that they may encode tail proteins involved in host range specificity.

**Figure 2.**
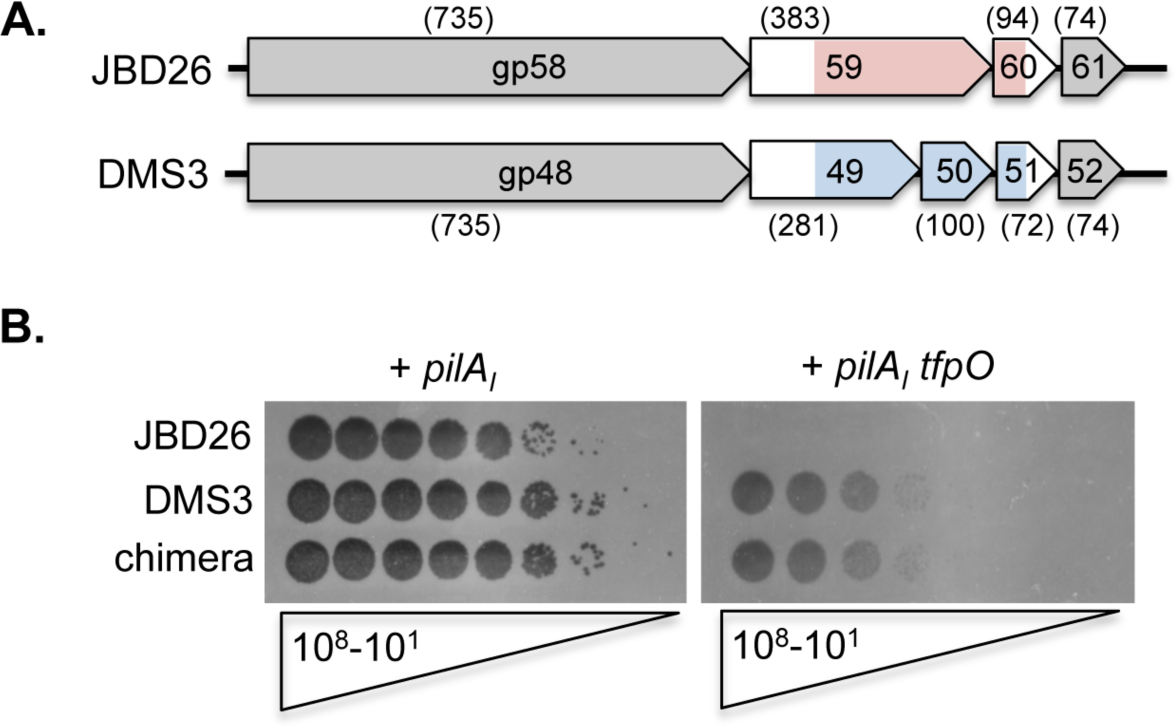
A chimeric phage expressing DMS3 genes in the JBD26 background gains the ability to infect a strain with glycosylated pilins. **A.** Map of JBD26 and DMS3 tail fibre genes and surrounding genomic conservation. The genomic organization of related genes from JBD26 and DMS3, and the predicted sizes of their products (amino acids, in brackets) are shown. Regions where sequences from the two phages diverge are colored pink and blue. The gp designations are omitted for most genes due to space limitations. The length of the predicted protein products in amino acids are shown in brackets above each open reading frame. Map not to scale. **B.** Plaque assay using 5 μl each of serial 10-fold dilutions of phage (10e8 – 10e1). A chimeric JBD26 phage expressing the DMS3 gp49-52 instead of its own gp59-60 gains the ability to infect a PAO1 *pilA* mutant complemented with the pilin from group I strain 1244 (*pilA_l_*), alone and with *tfpO*, encoding the cognate glycosyltransferase.

To determine if the differences in these putative tail proteins were responsible for unique ability of DMS3 to infect strains with glycosylated pilins, we generated a JBD26 lysogen of strain PAO1, then swapped some of its tail protein genes for those of DMS3 via homologous recombination. Open reading frames *59* and *60* of JBD26 were replaced with *49, 50* and *51* of DMS3, using the conserved flanking genes *58/48* and *61/52* as regions of homology for recombination (**Fig. 2A**). Strikingly, the resulting JBD26/DMS3 chimeric phage infected strains expressed both non-glycosylated and glycosylated pilins to the same extent as DSM3 (**Fig. 2B**). These results imply that the region of DMS3 encompassing ORFS *49-51* encodes the unique ability of DMS3 to use the glycosylated pilus for infection.

### Pilin sequence modulates phage susceptibility

The C-terminal Ser that is glycosylated in group I pilins is adjacent to the D-region of PilA ^23^, suggesting that post-translational modification could mask this potential phage binding surface (**Fig. 3A**). To test whether phages recognize the D-region of PilA, we exploited a set of previously engineered PilA_II_ single-residue substitution mutants ^34^. All mutants tested had similar levels of surface piliation (**Fig. 3B**), but plaquing assays revealed that a PilAII P138R mutant was approximately 1000-fold more resistant to JBD26 and JBD68 infection than wild type, while the effect of this mutation on susceptibility to DMS3 and the chimeric phage was minimal (**Fig. 3C**). A P138A substitution had no effect on susceptibility, while a P138M mutant had slightly reduced susceptibility, implying a specific interaction between phages JBD26 and JBD68 and the D-region of PilA_II_ that is disrupted by changes in side chain electrostatics. These results indicate that DMS3 and the JBD26 chimera containing DMS3 sequences interact with PilA in a different manner than JBD26 and JBD68, potentially explaining the ability of DMS3 to infect strains that express glycosylated subunits.

**Figure 3.**
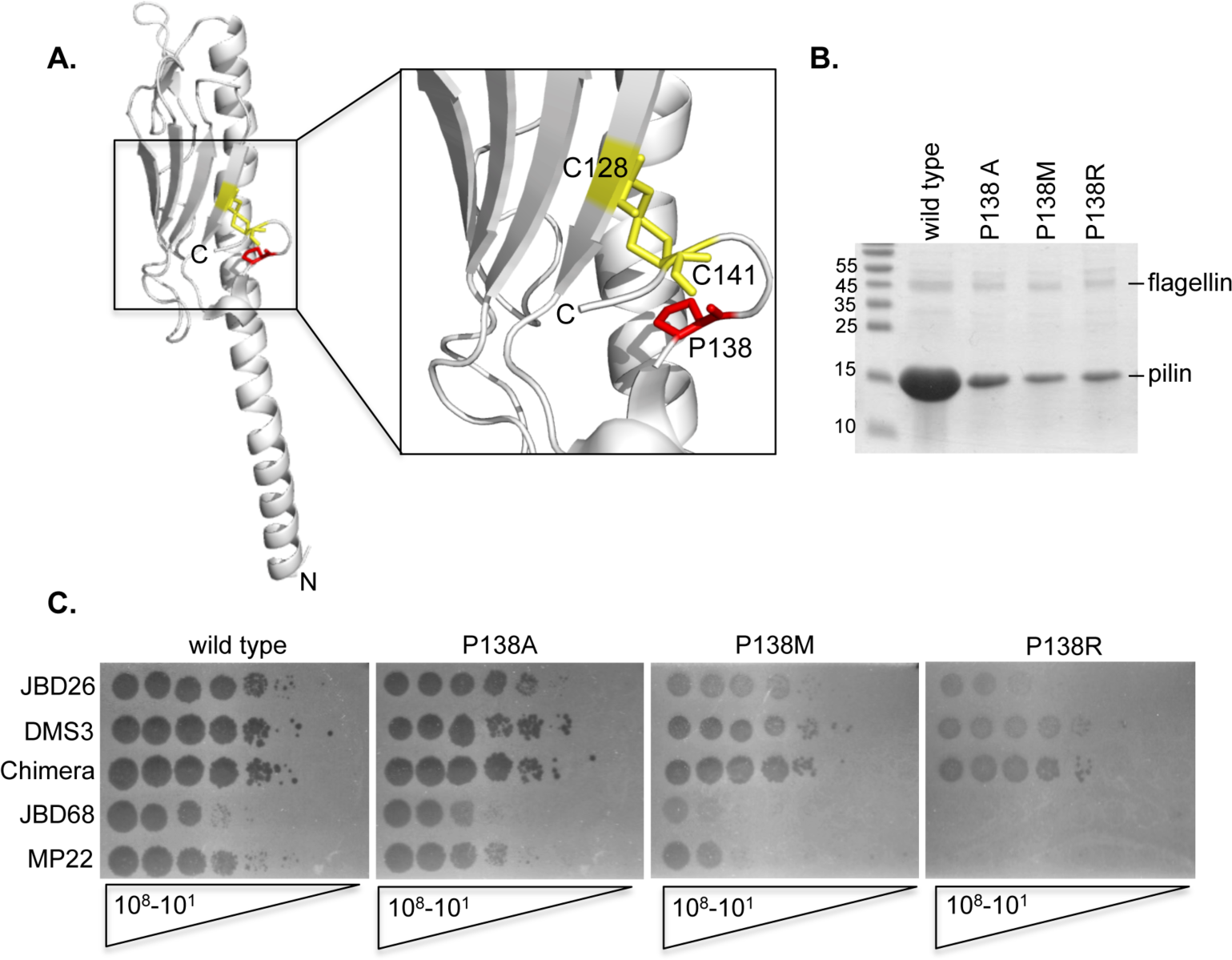
Pilin D-region sequence affects phage susceptibility. **A.** Model of a group II pilin showing the disulfide bonded Cys residues (yellow) demarking the C-terminal D region and the Pro138 residue (red) targeted for site directed mutagenesis. **B.** Relative surface piliation of the PilA_II_ point mutants. All three mutants have similar levels of piliation_34_. **C.** Plaque assays using 5 µl each of serial 10-fold dilutions of the phage indicated on the left (10e8 – 10e1). Expression of the P138R pilin in the PAO1 *pilA* background reduces or blocks infection by JBD26, JBD68, and MP22, but not DMS3 or the JBD26 chimera expressing DMS3 tail proteins, implying that DSM3 binds elsewhere.

### Pilin glycosylation with D-arabinofuranose also blocks phage infection

*P. aeruginosa* strains expressing group IV pilins – such as PA7 and Pa5196 – encode a distinct pilin glycosylation system that adds homooligomers of D-Ara*f* to multiple Ser and Thr residues on PilA_IV_^25^. To investigate whether this form of glycosylation also serves as a phage defense mechanism, we tested a set of PAO1 *pilA* mutant strains expressing unmodified or glycosylated PilA_IV_. Because JBD26 does not infect strains expressing PilA_IV_ well, we turned to a closely related phage, MP22^35^. MP22 gp54 has 99% identity to JBD26 gp59 that putatively recognizes unglycosylated pilins, and it has a similar infection pattern (**Fig. 3C**). We tested susceptibility of strains expressing D-Ara*f* glycans to MP22, JBD68, DMS3, and the chimeric phage (**Fig. 4**).

**Figure 4.**
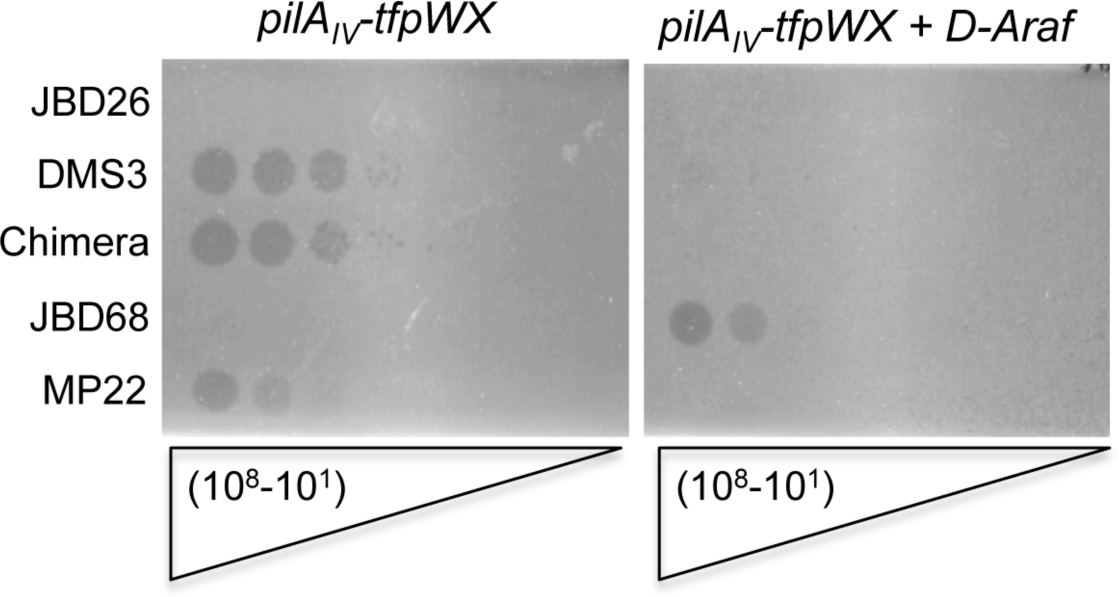
Pilin D-arabinofuranosylation blocks phage infection. Plaque assays using 5 µl each of serial 10-fold dilutions of the phage indicated on the left (10e8 – 10e1). Expression of PilA_IV_ and its accessory proteins TfpWX in the PAO1 *pilA* background results in expression of unmodified pilins (left). Addition of the genes encoding the synthesis of D-arabinofuranose polymers (*D-Araf*) from pUCP20-6245-51 results in expression of glycosylated pilins^26^and blocks infection by DMS3, the chimeric JBD26 phage, and MP22.

Unlike glycosylation of pilins with O-antigen units, this form of glycosylation prevented infection by DSM3 and the chimera, as well as MP22. Interestingly, JBD68 could not infect the strain that produced unmodified PilA_IV_, but could infect the strain expressing glycosylated PilA_IV_, suggesting that it recognizes a new epitope present on the modified pilin, or the unique pilin glycans themselves. These data show that both of *P. aeruginosa’s* pilin glycosylation systems have the capacity to protect against subsets of pilus-specific phages. In addition, phages like JBD68 appear to have evolved to exploit some forms of pilin glycosylation for cell surface binding.

## DISCUSSION

Since glycosylation of thousands of surface-exposed pilin subunits can be energetically costly (each sugar requires the production of a nucleotide-activated precursor, and each glycan unit contains multiple sugars), the addition of such modifications must provide a valuable benefit. Here, we showed that pilin glycosylation provides protection against many pilus-specific bacteriophages. Given the powerful impact of phage predation on bacterial evolution, our data provide a compelling rationale for the widespread glycosylation of pilins in environmental and pathogenic bacteria. Glycosylation of *P. aeruginosa* pilins was previously suggested to protect against complement binding, opsonization by pulmonary surfactant protein A, or against proteolytic degradation by host or self-produced proteases ^36, 37^. Similar hypotheses have been advanced to explain the function of pilin glycosylation in *Neisseria* and *Acinetobacter* ^31, 32^. While these effects of pilus glycosylation may have some impact within the context of infection, resistance to phage predation is likely of greater advantage to bacteria in a much broader range of native environments and conditions.

A recent study of *P. aeruginosa* phages from cystic fibrosis sputum samples suggested that phage density was ~10-100 fold greater than that of *P. aeruginosa*, and that lytic phage activity controls bacterial numbers in chronically infected patients ^38^. Pilin glycosylation thus has the potential to influence virulence both directly – blocking antibody and complement binding – and indirectly, allowing bacteria to make functional pili and maintain high bacterial loads in the presence of pilus-specific phages. These features may account for the reported prevalence (~70%) of group I strains of *P. aeruginosa* among CF isolates ^22^, as such strains are likely to be more generally resistant to phage infection than strains producing non-glycosylated pili. The phage protection afforded by pilin glycosylation may also explain why group I and group IV pilins assemble poorly in the absence of this modification (**Fig. 1**)^26^. Limiting the assembly of unmodified pilins may be a quality-control mechanism that reduces the dangerous possibility that non-glycosylated – and thus phage-vulnerable – pili are expressed on the cell surface. Although loss of pilus expression or function can also protect the bacteria from phage predation, these are disadvantageous phenotypes that reduce fitness ^39, 40, 41^.

While T4P are well-established receptors for phage, the exact mode of phage binding has not been clarified. Electron micrographs show binding of phage along the length of pilus filaments ^42^, presumably to the major pilin. Our data strongly support the conclusion that phages bind directly to PilA, rather than any of the minor pilins, for the following reasons: 1) Glycosylation of PilA blocks phage infection, and the minor pilins are not glycosylated ^18, 20, 43^. 2) The plaquing ability of all the phages tested was affected by the group and/or glycosylation state of the major pilin, and all the assays were performed in the PAO1 background where the minor pilins were invariant. 3) Specific single amino acid substitutions in PilA changed the susceptibility of strains to phage infection (**Fig. 3**). Although we cannot rule out that some pilus-specific phages may bind to minor pilins, the phages studied here interact with the major pilin. Given that minor pilins are mainly localized to the pilus tip and are present in minute quantities relative to the major pilin, the binding of phages to PilA is expected to be a superior strategy for cell surface engagement.

The ability of the phages used in this study to use both unmodified PilA_I_ and PilA_II_ as receptors is remarkable, as these proteins are only 25% identical outside of the highly conserved N-terminal 25 residues that are largely buried in the interior of assembled pilus filaments ^44, 45^. The binding of phages to diverse PilA proteins likely reflects the results of the evolutionary drive for phage to use a wide variety of pili as receptors, and further emphasizes the difficulty of evading phage predation only through accumulation of amino acid substitutions in PilA, which could also have negative functional consequences. Pilus glycosylation may provide a more effective defense against phages, by masking phage-binding sites through steric hindrance. Interestingly, some pilus-specific phages such as JBD68 appear to have evolved the capacity to recognize pilins with specific post-translational modifications (**Fig. 4**), potentially broadening their host range.

A role for DMS3 gp49, gp50 and/or gp51 in recognizing the glycosylated pilus was demonstrated by recombining the genes encoding these proteins into JBD26, allowing this phage to use the glycosylated group I pilus for infection. DMS3 gp49 and JBD26 gp59 each have a conserved 100 residue N-terminal domain that corresponds to the Pfam protein family DUF2793 (PF10983). Proteins containing this domain are encoded in a variety of non-contractile tailed phage and prophage genomes, with the DUF2793 protein invariably encoded immediately downstream of the gene encoding the central fibre protein ^46^ as in JBD26 and DMS3 (**Fig. 2A**). Central fibre proteins and others encoded in the 3’-regions of tail operons are often involved in host cell receptor binding ^46, 47^. DUF2793 proteins are variable in length and display highly divergent C-terminal regions. The conserved N-termini likely attach these proteins to the phage particle while the C-termini may interact with the variable surfaces of host cells. Among the functions predicted for the variable C-terminal domains are peptidase and glycanase activities (**Supplementary Fig. S1B**). This modular organization may prove useful for future development of designer phages that can target specific species or strains. Mass spectrometry on purified phages closely related to JBD26 and DMS3 (**Supplementary Table S1**) confirmed that the DUF2793 proteins are present in phage particles. We conclude that gp49 and gp59 are the components of DMS3 and JBD26 tails, respectively, that are responsible for binding to PilA. The difference in the effects of PilA mutations on infectivity of JBD26 versus DMS3 (**Fig. 3C**) implies that DMS3 gp49 and JBD26 gp59 interact with different regions of PilA, accounting for DMS3’s ability to bind glycosylated PilAI, while JBD26 gp59 cannot. In contrast, the inability of DMS3 gp49 to bind glycosylated PilAIV is likely due to the different position of these modifications on the protein ^25^. The identical plating behavior of the JBD26/DMS3 chimeric phage and DMS3 implies that gp49 is responsible for conferring the unique host specificity of DMS3. The functions of DMS3 gp50 and gp51 are not known, and homologues of gp51 were not found in phage particles (**Supplementary Table S2**).

The function of type IV pilin glycosylation has long remained a mystery. We show here that both of *P. aeruginosa*’s glycosylation systems block infection by a variety of pilus-specific phages, and suggest that resistance to phage attack is the major evolutionary driving force for these post-translational modifications. Since pili are common targets for phage adsorption in many species, pilus glycosylation may be a widespread anti-phage defense system. The large size of oligosaccharide groups and their typically negative charge can serve to effectively block phage accessibility to large portions of the pilus surface ^31^, and thus engender resistance to diverse phages. In this way, glycosylation is a more broadly useful resistance mechanism than amino acid substitutions in PilA, a conclusion that is supported by the ability of the phages tested here to utilize PilA proteins of diverse sequence as receptors. Glycosylation also provides a means to block phage infection without compromising the important adaptive functions of the type IV pilus. In conclusion, pilus glycosylation is another fascinating manifestation of the evolutionary battle between bacteria and phages. The ability of phages DMS3 and JBD68 to each partially overcome specific forms of pilus glycosylation is likely a foreshadowing of a variety of mechanisms by which phages can circumvent this defense that await discovery.

## METHODS

### Bacterial strains, phage, and plasmids used in this work

Bacterial strains, phage, and plasmids are listed in **Supplementary Table S3**. Unless noted otherwise, *E. coli* and *P. aeruginosa* were grown at 37°C in Luria-Bertani (LB) broth or 1.5% agar plates supplemented with antibiotics at the following final concentrations when necessary (μg/mL): ampicillin (Ap), 100; kanamycin (Kn), 50; gentamicin (Gm), 15 for *E. coli* and 30 for *P. aeruginosa*. Plasmids were transformed by heat shock into chemically competent *E. coli* cells or by electroporation or biparental mating with *E. coli* SM10 into *P. aeruginosa* as previously described ^26^. All constructs were verified by DNA sequencing.

### Phage plaquing assays

Phage plaque assays were performed as described previously ^48^, with modifications. Briefly, bacteria were grown overnight at 37°C with shaking, and then subcultured 1:100 in LB plus 8 mM MgSO4 and grown for 3 h at 37°C with shaking. The subculture was then standardized to OD_600_ of 0.3 in LB plus 8 mM MgSO4 and 100 μl was mixed with 8 ml top agar (LB plus 8 mM MgSO4, 0.6% agar), which was overlaid on a pre-poured rectangular LB plus 8 mM MgSO4 1.5% agar plate containing antibiotics and L-arabinose where indicated. After allowing the top agar to solidify, it was air-dried with the lid off in a biosafety cabinet for 15 min. Phage stocks were standardized to a PFU/ml of 10e8 and 10-fold serially diluted in LB plus 8 mM MgSO4, and 5 μl of each dilution was spotted onto the prepared plates. After allowing the spots to air dry for 10 min with lid on, the plates were incubated inverted for 18 h at 37°C. The plates were then photographed and the titre estimated as the lowest dilution generating a complete zone of cell lysis.

### Twitching motility assays

Twitching assays were performed as described previously ^29^. Single colonies were stab inoculated to the bottom of a 1% LB agar plate which was incubated for 36 h at 37°C. The agar was then carefully discarded, and the adherent bacteria stained with 1% (w/v) crystal violet dye, followed by washing with tap water to remove unbound dye. Twitching zone areas were measured using ImageJ software (NIH). All experiments were performed in triplicate with at least three independent replicates.

### Sheared surface protein preparation

The levels of surface-exposed pili and flagella were analyzed as described previously ^26^. Briefly, the strains of interest were streaked in a grid-like pattern on LB agar plates and incubated at 37°C for ~16 h. The cells were scraped from the plates with glass coverslips and resuspended in 4.5 mL of phosphate buffered saline, pH 7.0. Surface proteins were sheared by vortexing the cell suspensions for 30 s. The suspensions were transferred to three separate 1.5 mL Eppendorf tubes and cells pelleted by centrifugation at 11,688 x *g* for 5 min. The supernatant was transferred to fresh tubes and centrifuged at 11,688 x *g* for 20 min to pellet remaining cells. After transfer of supernatants to new tubes, the surface proteins were precipitated by adding 1/10 volume of 5M NaCl and 30% (w/v) polyethylene glycol (PEG 8000, Sigma Aldrich) to each tube and incubating on ice for 90 min. Precipitated proteins were collected by centrifugation at 11,688 x *g*, and resuspended in 150 μL of 1X SDS sample buffer (125mM Tris, pH 6.8, 2% β-mercaptoethanol, 20% glycerol, 4% SDS and 0.001% bromophenol blue). Samples were boiled for 10 min and separated on 15% SDS-PAGE gels. Proteins were visualized by staining with Coomassie brilliant blue.

### Generation of a JBD26-DMS3 chimeric phage

To create a JBD26 chimeric phage expressing DMS3 tail protein, we first generated a JBD26 lysogen of strain PAO1. The primers chimera1 5’- ACAAGA<u>GAATTCC</u>AGGATCAGTTCAATCTG-3’ and chimera2 5’- ACAAGA<u>CTGCAG</u>CTAGCCATTGTGCTGTAGCG-3’, containing EcoRI and PstI restriction sites (underlined), were used to amplify from the middle of DMS3 gp48 to the end of gp52 (~3 kb). The resulting PCR product was gel-purified and ligated into the suicide vector, pEX18Gm ^49^. After validation of the construct by DNA sequencing, it was introduced into *E. coli* SM10 by CaCl_2_ transformation and transferred to the PAO1 JBD26 lysogen by biparental mating as described previously ^26^. Following overnight incubation at 37°C, mating mixtures were resuspended in 1 ml LB and 100 μl was plated onto *Pseudomonas* Isolation agar (PIA) containing 100 μg/ml of Gm to counterselect the *E. coli* donor. Colonies were picked from PIA onto LB Gm30 agar and from there, onto LB no salt, 5% sucrose agar to select against merodiploids. Colonies that grew on sucrose plates were streaked in parallel on LB Gm30 and LB agar, and Gm sensitive colonies were tested by colony PCR using primers DMS3gp50p1 5’- GAACAGAATTCGAGGTGGTTCTGATGATGATCATC-3’ DMS3gp50p2 5’- GAACAGGATCCTCATTGCGGCAACTCCACAGG-3’, designed to amplify DMS3 gp50. DNA from PCR-positive colonies was reamplified using primers DMS3gp49p1 5’- GAACAGAATTCAGGAGGCGTATCCGCATGAGC-3’ and DMS3gp52p2 5’- ACAAGACTGCAGCTAGCCATTGTGCTGTAGCG-3’, and the resulting amplicons sequenced to verify gene replacement.

To induce the excision of the chimeric phage, the lysogen was grown overnight in LB and subcultured 1:100 to OD_600_ 0.6 in 3 ml of the same medium, then treated with mitocycin C (final concentration 3 μg/ml) for 18 h. The remaining cells were lysed by adding ~250 μl chloroform and vortexing briefly to mix. The phage were titred by standard plaque assay. Ten-fold serial dilutions of the lysate were prepared and 100 μl each mixed with 100 μl of PAO1 standardized to OD600 0.3 in LB plus 10 mM MgSO4 and added to 8 ml of top agar (LB plus MgSO4, 0.6% agar) that was overlaid onto a standard LB plus 10 mM MgSO4 plate and allowed to solidify. After 18h incubation at 37C, the dilution(s) giving countable plaques were used to calculate the plaque-forming units per ml of lysate.

### Mass spectrometry analysis of phage particles

We subjected 3.8x10^9^ phage particles from lysate purified twice by CsCl density gradient ultracentrifugation to tryptic digest as described ^50^. LC-MS/MS spectra were collected on a linear ion-trap instrument (ThermoFisher LTQ)(*SPARC* BioCentre, The Hospital for Sick Children, Toronto, Canada). Proteins were identified using Mascot (Matrix Science, London, UK) and analyzed in Scaffold version 3.0 (Proteome Software Inc., Portland, OR, USA). The protein identification cut off was set at a confidence level of 95% with a requirement for at least two peptides to match to a protein.

## ACKNOWLEDGEMENTS

We thank Karen Maxwell for helpful comments on the manuscript. This work was supported by Canadian Institutes of Health Research (CIHR) Open Operating Grants to LLB (MOP 86639) and to ARD (XNE-86943 and MOP-115039).

## AUTHOR CONTRIBUTIONS

HH, ARD, and LLB designed the study; HH, JBD, HM, and KMS performed experiments; HH, ARD, and LLB analyzed the data; ARD and LLB wrote the manuscript with input from JBD and HM. All authors approved the final version.

